# The *Medicago truncatula* Vacuolar Iron Transporter-Like proteins VTL4 and VTL8 deliver iron to endosymbiotic bacteria at different stages of the infection process

**DOI:** 10.1101/689224

**Authors:** Jennifer H. Walton, Gyöngyi Kontra-Kováts, Robert T. Green, Ágota Domonkos, Beatrix Horváth, Ella M. Brear, Marina Franceschetti, Péter Kaló, Janneke Balk

## Abstract

The symbiotic relationship between legumes and rhizobium bacteria in root nodules has a high demand for iron. The host plant is known to provide iron to developing bacteroids, but questions remain regarding which transporters are involved. Here, we characterize two Vacuolar Iron Transporter-Like (VTL) proteins in *Medicago truncatula* that are specifically expressed during nodule development. *VTL4* is mostly expressed during early infection and the protein was localized to membranes and the infection thread. *vtl4* mutants were delayed in nodule development. *VTL8* is closely related to *SEN1* in *Lotus japonicus* and expressed in the late stages of bacteroid differentiation. The VTL8 protein was localized to the symbiosome membrane. A mutant line lacking the tandemly-arranged *VTL4* – *VTL8* genes, named 13U, was unable to develop functional nodules and failed to fix nitrogen, which was restored by expression of *VTL8* alone. Using a newly developed *lux* reporter to monitor iron status of the bacteroids, a slight decrease in luminescence was observed in *vtl4* mutants and a strong decrease in the 13U mutant. Iron transport capability of VTL4 and VTL8 was shown by yeast complementation. Taken together, these data indicate that VTL-type transporters are the main route for delivering iron to symbiotic rhizobia.

## INTRODUCTION

Legumes and a small number of other plant species (*Parasponia* sp.) are able to form a symbiosis with rhizobium bacteria which enables the host plant to access N_2_ as a source of nitrogen. The host plant provides carbohydrates derived from photosynthesis for the energy-demanding reduction of N_2_ to ammonium, carried out by the bacterial nitrogenase enzyme. Successful establishment of the symbiosis in specialized structures called root nodules depends on signalling and recognition between the rhizobia and the host plant, as well as later checkpoints during nodule development (reviewed in Oldroyd *et al*., 2011; Suzaki *et al*., 2015).

Root nodules have a high requirement for iron owing to abundant iron proteins. Infected plant cells produce large amounts of haem as a cofactor in leghaemoglobins which maintain a microaerobic environment for the oxygen-sensitive nitrogenase enzyme (Downie, 2005; Ott *et al.*, 2005; Garrocho-Villegas *et al.*, 2007). The bacterial nitrogenase enzyme binds 12 iron in the form of iron-sulphur clusters and another 7 in the Fe-Mo cofactor (Dean *et al.*, 1993; Howard & Rees, 2006). The nitrogen-fixing bacteria also contain numerous cytochromes and other iron proteins. When the plant is starved of iron, nodule initiation and further development is strongly impaired (O’Hara *et al.*, 1988; Tang *et al.*, 1990; Brear *et al.*, 2013).

Nodule development is initiated by a signalling cascade between plant roots and the bacteria, entrapment of the bacteria in curled root hairs and formation of infection threads along which the bacteria travel into the root cortex. Division of cortex cells leads to formation of a specialized root outgrowth, the nodule. The root nodules formed by *Medicago truncatula* in symbiosis with *Sinorhizobium meliloti* or *S. medicae* are of the indeterminate type. The meristem persists over time and generates a gradient of cells at progressing developmental stages (Vasse *et al.*, 1990; Roux *et al.*, 2014), commonly divided into 4 histological zones, with the meristem as Zone I. Zone II corresponds to the infection zone where bacteria are released from the infection threads, divide and form symbiosomes – intracellular bacteria surrounded by a plant-derived membrane. In older cells of Zone II bacterial cell division stops while genome replication continues, resulting in elongated polyploid bacteroids (Mergaert *et al.*, 2006). Synchrotron X-ray fluorescence (XRF) studies on *M. truncatula* indicated that iron is delivered to Zone II by the vascular bundles in the nodule (Rodríguez-Haas *et al.*, 2013). The expression of the iron storage protein ferritin is induced in Zone II (Strozycki *et al.*, 2007; Roux *et al.*, 2014), corresponding to ‘iron-rich bodies’ in and near the vascular bundles detected by XRF (Rodríguez-Haas *et al.*, 2013). In the Interzone (transition of zone II-III), infected plant cells and the bacteroids complete their differentiation with the last rounds of endoreduplication and enlargement, resulting in a striking cell morphology of tightly packed elongated rhizobia surrounding a central vacuole. Zone III comprises the major part of a functional nodule and this is where nitrogen-fixation takes place. In older nodules, a zone of senescent cells, Zone IV, is formed from which nutrients such as iron are recycled from degraded plant cells and symbionts (Van de Velde *et al.*, 2006; Rodríguez-Haas *et al.*, 2013).

Forward and reverse genetic studies have identified several transporters involved in iron delivery to the nodules and symbiosomes. The closely related genes *L. japonicus MATE1* and *M. truncatula MATE67* are highly induced during nodule development, specifically in the infection zone (Takanashi *et al.*, 2013; Wang *et al.*, 2017; Kryvoruchko *et al.*, 2018). Studies in *Xenopus* oocytes showed that the LjMATE1 and MtMATE67 proteins can transport citrate, an organic chelator of Fe^3+^ in the xylem (Takanashi *et al.*, 2013; Kryvoruchko *et al.*, 2018). The MtMATE67 protein is localized to vascular bundles but also to the symbiosome membrane (Kryvoruchko *et al.*, 2018). An RNAi mutant of *LjMATE1* accumulated high levels of iron in the vascular bundle of roots and nodules, whereas the iron concentration in infected cells of the nodules was lower compared to wild type (Takanashi *et al.*, 2013). These two studies indicate that the host plant delivers iron to the nodules via the xylem.

Uptake of iron into the infected cells may involve *MtNRAMP1*, one of 7 members of this gene family in *M. truncatula* with the highest relative expression in roots and nodules (Tejada-Jiménez *et al.*, 2015). Yeast complementation studies showed that MtNRAMP1 can transport iron and manganese, similar to its closest Arabidopsis homologue which is the primary route for manganese into the cell (Cailliatte *et al.*, 2010). MtNRAMP1 is localized to plasma membranes, including the plasma membrane of infected cells. A transposon insertion mutant of *MtNRAMP1* has impaired nodule development, but still exhibited 60% nitrogenase activity compared to wild type (Tejada-Jiménez *et al.*, 2015).

For iron to reach the bacteroid, it needs to be exported by the infected plant cell and imported across the bacteroid membrane. Five different allelic mutants in the *LjSEN1* gene, encoding a Vacuolar iron Transporter-Like (VTL) protein, were identified in a screen for ineffective symbiotic (Fix-) mutants (Hakoyama *et al.*, 2011). The VTL proteins differ from the better characterized Vacuolar Iron Transporter (VIT) proteins in that they lack a cytosolic loop thought to mediate Fe^2+^/H^+^ antiport (Labarbuta *et al.*, 2017; Kato *et al.*, 2019). Promoter-GUS studies showed that *LjSEN1* is expressed exclusively in rhizobia-infected cells, but its subcellular localization and function in iron transport has not been documented to date.

To better understand the role of VTLs in nodule development and biological nitrogen fixation, we characterized two *VTLs* in *M. truncatula, VTL4* and *VTL8*, which are exclusively expressed in nodules. Localization and mutant studies showed they play a role at early and late stages, respectively, of nodule development. VTL8 is critical for delivering iron to the bacteroids and establishment of the nitrogen-fixing bacteria.

## RESULTS

### *M. truncatula* has 8 *VTL* genes of which two genes are expressed in nodules

The *M. truncatula* genome (Mt4.0v1) was searched for homologs of *A. thaliana VIT* and *VTL* genes. We identified 2 homologs of *VIT* and 8 homologs of *VTL* as compared to one and 5 genes, respectively, in *A. thaliana*. Amino acid alignment of the 8 MtVTLs with the *A. thaliana* homologs showed only a limited phylogenetic relationship between different VTLs of the two species (Fig. 1a). Based on this phylogeny, the *M. truncatula* genes were assigned as *VTL1-VTL8. MtVTL1, MtVTL2* and *MtVTL3* are most similar to *A. thaliana VTL1–VTL4. MtVTL4, MtVTL5, MtVTL6* and *MtVTL7* are closely related paralogs on chromosome 4 which are more similar to *A. thaliana VTL5* than the other *AtVTL* genes. In contrast, *MtVTL8* has no direct homologue in *A. thaliana*, but is closely related to *NODULIN-21* of soybean (Delauney *et al.*, 1990) and *SEN1* in *L. japonicus* (Hakoyama *et al.*, 2011), which have strongly induced expression in nodules. Indeed, *MtVTL8* is also highly expressed during the later stages of nodule development, with the highest expression in the Interzone (Zone II – III), see Fig. 1b based on data in Roux *et al*., (2014). Of the other *MtVTL* genes, only *MtVTL4* transcripts are found at significant levels in nodules, with the highest levels in the early infection stage (distal zone II, ZIId).

**Fig. 1.**
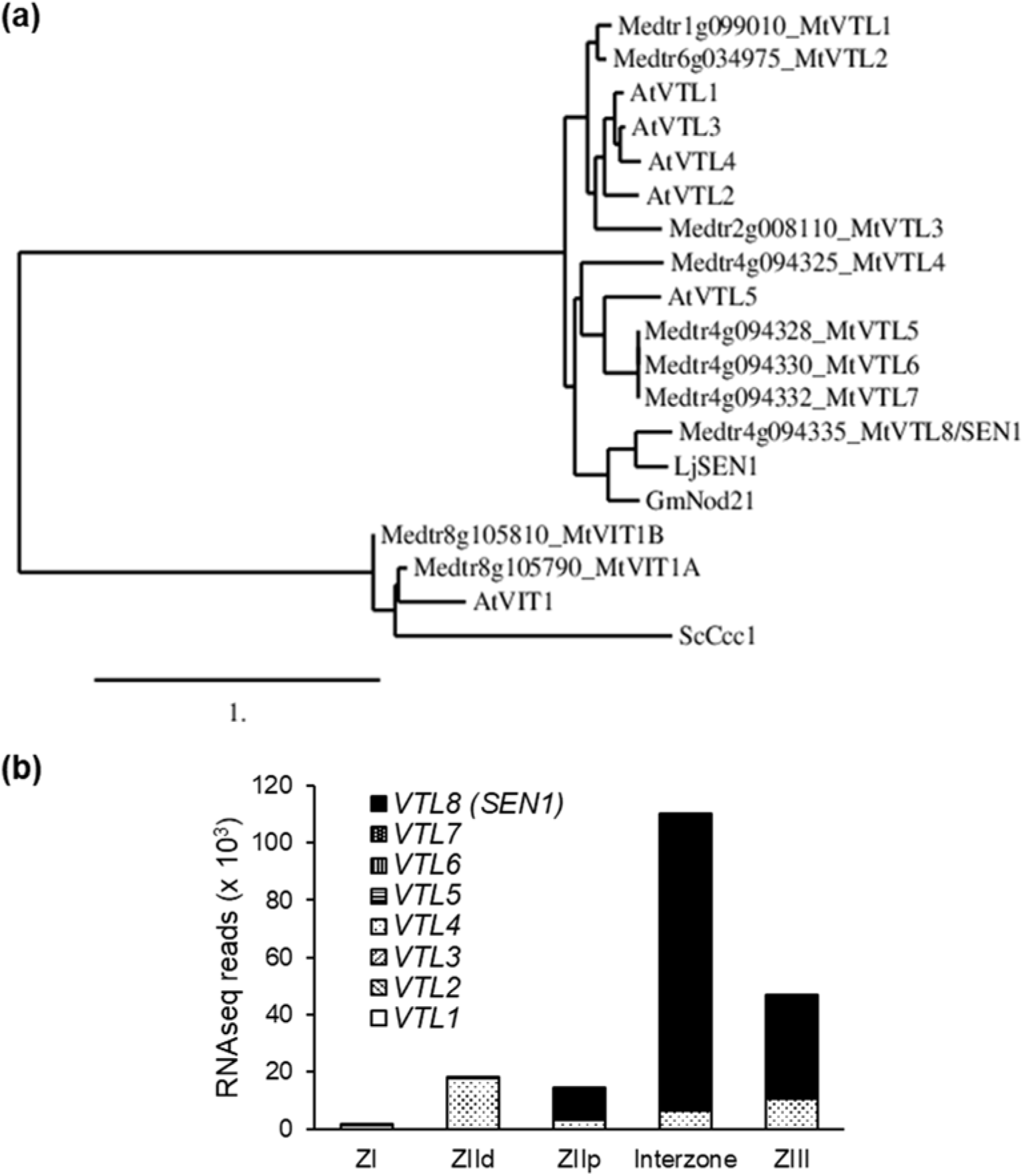
Vacuolar iron Transporter-Like (VTL) proteins in *Medicago truncatula*. (a) Phylogenetic relationship of VTL protein sequences identified in the *M. truncatula* genome and those previously reported in *Arabidopsis thaliana* (*AtVTLI, AT1G21140*);, *AtVTL2, AT1G76800; AtVTL3, AT3G43630; AtVTL4, At3G43660; AtVTL5, AT3G25190).* The ‘true’ Vacuolar Iron Transporter (VIT) sequences including Ccc1 from *Saccharvmyces cerevisiae* served as an outgroup. The phylogenetic tree was generated using the server www.phylogeny.fr (b) Expression levels of *VTL* genes in different nodule zones. Data were obtained from RNAseq data published by Roux *et al.*, (2014).

### MtVTL4 and MtVTL8 localize to membranes in different zones of nodule development

To investigate the localization of the VTL4 and VTL8 proteins in nodules, the proteins were fused via a short C-terminal linker peptide to mCherry and transiently expressed in roots under the control of their own promoters. The presence of a glycine-rich linker peptide has previously been shown to improve membrane insertion and thus stability of *A. thaliana* VTLs expressed in Baker’s yeast (Gollhofer *et al.*, 2014). As a marker protein for the plasma membrane we used the aquaporin PIP2A from *A. thaliana* (Cutler *et al.*, 2002) which localizes similarly to the cell membrane in *M. truncatula* roots (Ivanov & Harrison, 2014). The symbiosome membrane is derived from the plasma membrane, however regulatory elements that target proteins there are not well understood. The coding sequence of *AtPIP2A* was fused to enhanced GFP and the resulting PM-eGFP marker was expressed using the *LjUBI* promoter. From several membrane markers that were tested, *LjUBI:AtPIP2A-eGFP* was expressed in all nodule cell types and GFP was found in the plasma membrane as well as the symbiosome membrane (Fig. 2).

**Fig. 2.**
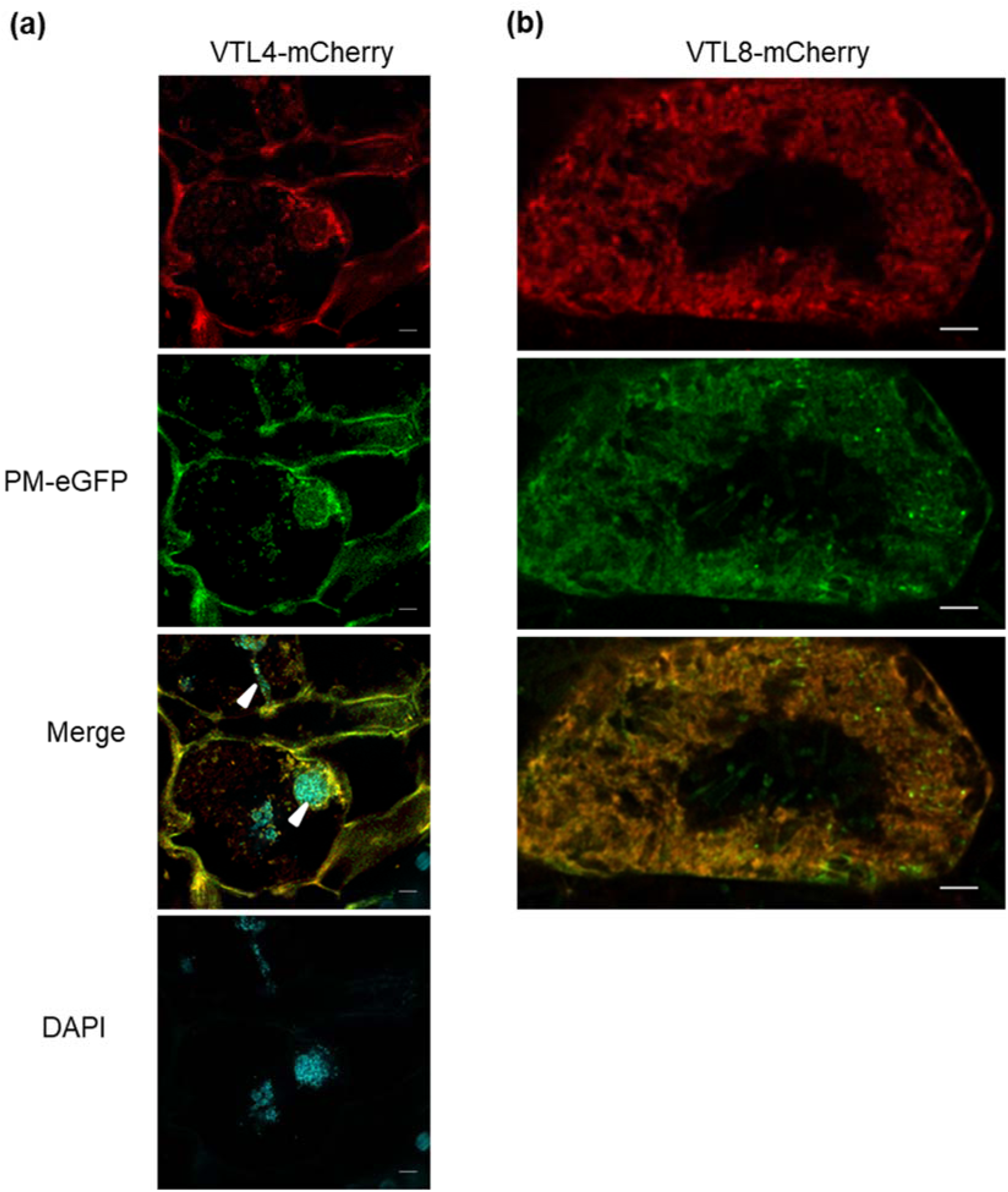
Localization of *VTL4* and *VTL8* in root nodules. (a) VTL4 fused to mCherry in cells of zone II (infection zone). The fusion protein was expressed using the *VTL4* promoter. The Arabidopsis *PIP2A* promoter and coding sequence fused to eGFP serves as a marker protein for plasma membrane (PM). DAPI staining visualizes DNA including the bacteria in infection threads (arrow heads). Scale bar is 5 μm. (b) Localization of VTL8 fused to mCherry in infected cells of the interzone (zone ll-lll). The fusion protein was expressed using the *VTL8* promoter. Scale bar is 5 μm.

In nodules expressing VTL4-mCherry, red fluorescence was observed in cells of the infection zone only, corresponding to the highest level of RNA expression in this zone (Fig. 1b). VTL4-mCherry co-localized with the PM-eGFP marker to plasma membranes. VTL4-mCherry was also associated with membranes of the infection thread (Fig. 2a, arrow head in merged image). VTL8-mCherry was detected in the infected cells of the interzone only (Fig. 2b). The fusion protein was confined to the bacteroids surrounding the central vacuole (upper panel). Co-localization with the PM-eGFP marker indicates that VTL8 is localized to the symbiosome membrane.

### *VTL8* is required for nodule development and nitrogen fixation

To study the role of *VTL4* and *VTL8* in nodule development, mutants were obtained from different sources. For *VTL4*, two lines with a *Tnt1* insertion in the coding sequence were isolated (Fig. 3a), but none were found for *VTL8* in the Noble Foundation collection despite extensive screening by PCR. However, a mutant line lacking several *VTL* genes was isolated from a collection of Fix-mutants generated by fast neutron radiation. The rough map position of the 13U mutant was identified previously between the genetic markers h2_32m20c and DENP (Domonkos *et al.*, 2013). Microarray hybridization using the genomic DNA of 13U identified a probe set corresponding to the gene *Medtr4g094325* encoding *VTL4*. Further analysis by PCR amplifications identified a 30-kb deletion in the 13U genome spanning *VTL4* to *VTL8*. Gene expression analysis confirmed the absence of *VTL4* transcript in the *vtl4* mutants and in the 13U line (Fig. 3b, c). While low levels of PCR product for *VTL8* were detected in the 13U mutant using standard RT-PCR (Fig. 3b), no product was generated by quantitative RT-qPCR (Fig. 3c). Possibly, aspecific priming to *VTL* paralogues occurred in the RT-PCR reaction, but not with more stringent qPCR conditions.

**Fig. 3.**
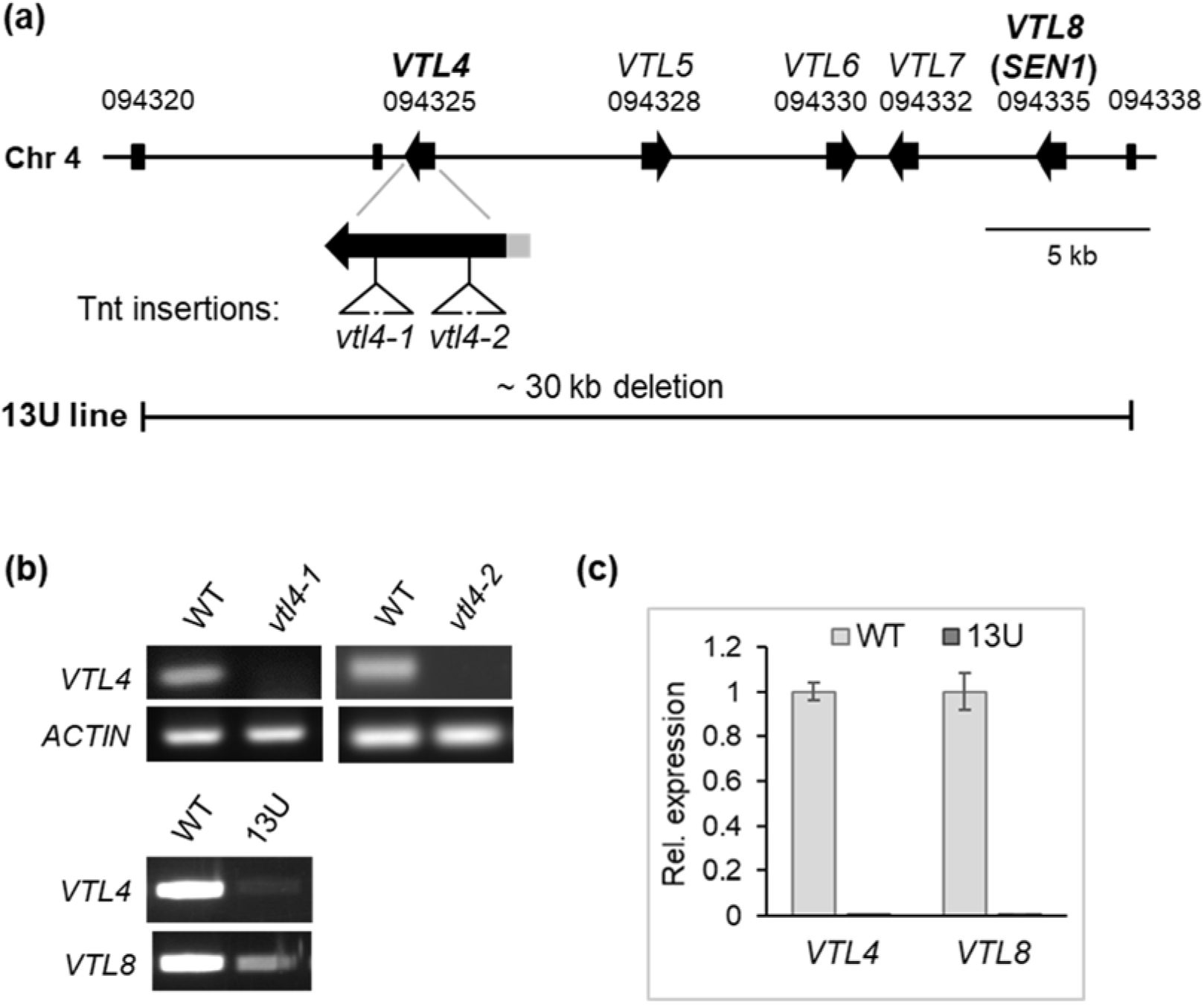
*M. truncatula* mutant lines affected in *VTL* gene expression. (a) Arrangement of *VTL* paralogs on chromosome 4. The position of two different *Tnt1* insertions in *VTL4* are indicated and also the deleted segment in the 13U line. (b) *VTL4* transcript levels in nodules (28 dpi) of the parental wild type (WT), the *vtl4-1* and *vtl4-2 Tnt1* insertion lines, NF17463 and NF21016, respectively, and the 13U line. (c) Expression of *VTL4* and *VTL8* in nodules of the 13U line relative to the parental wild type line (WT), at 28 dpi. as determined by RT-qPCR. The expression was normalized to that of the *UPL7* gene.

13U plants grew less vigorously than the parental wild type (Fig. 4a) and nodules were arrested early in development, lacking the typical pink colour of leghaemoglobin (Fig. 4b). Previous studies showed bacterial colonization of the infection zone and interzone of the 13U mutant nodules and only sporadic infection within the nitrogen fixation zone (Domonkos *et al.*, 2013; Fig. 4c,d). To investigate the morphology and arrangement of rhizobia, nodule sections were stained with the nucleic acid binding dye SYTO13. No significant difference was found in the morphology of bacteria, but in the differentiation zone of 13U mutant nodules the elongated rhizobia were disorganized rather than orientated toward the vacuoles of infected cells found in wild type (Fig. S1).

**Fig. 4.**
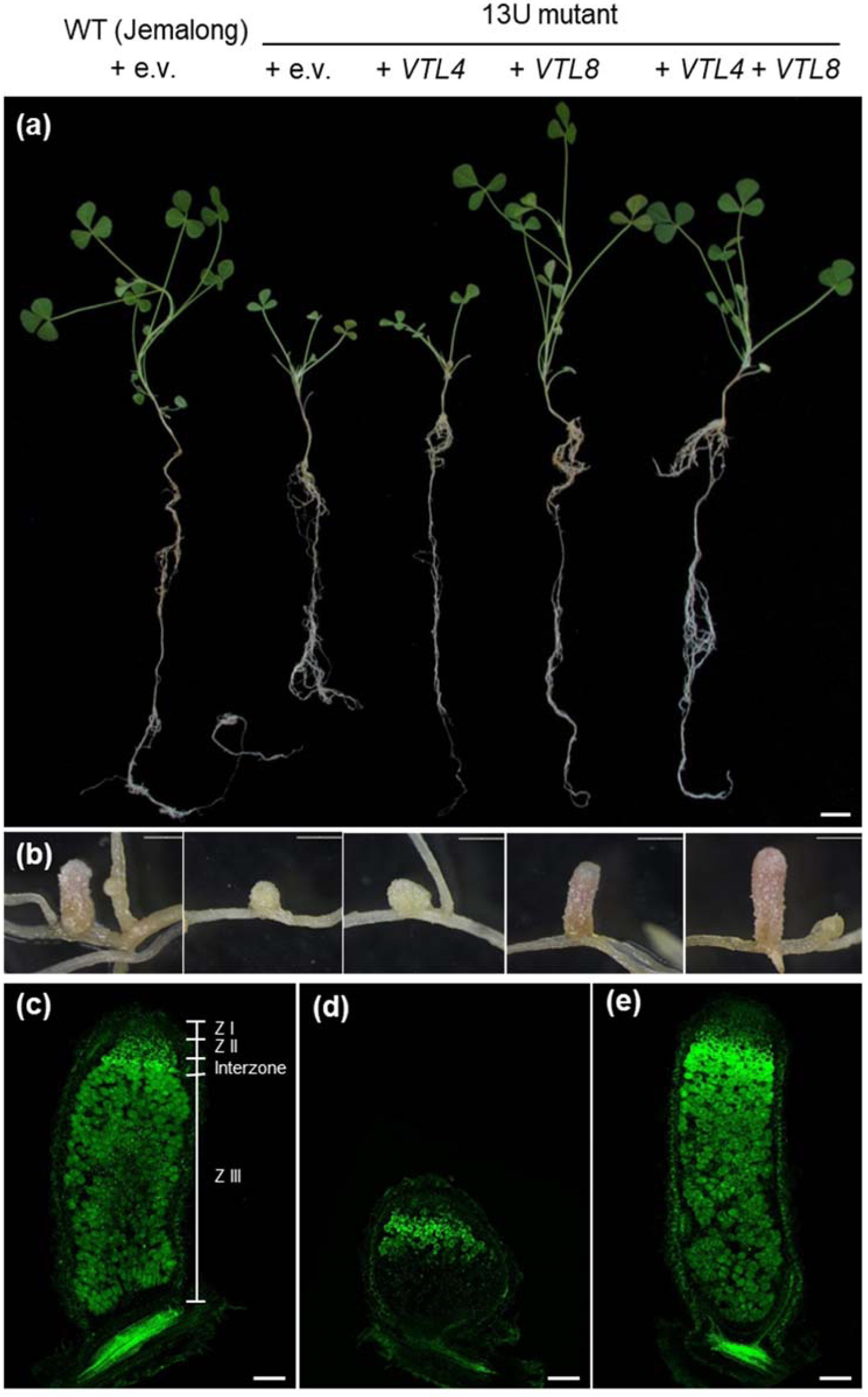
*VTL8* is required for development of functional nodules. (a) Wild type (WT Jemalong) plants and the 13U mutant transformed with *VTL4, VTL8, VTL4* + *VTL8* or empty vector (e.v.), as indicated. Roots were inoculated with *S. medicae* and images taken 31 dpi. Scale bar is 1 cm. (b) Root nodules from plants in (a). Transformed roots were confirmed by DsRed fluorescent protein marker (not shown). Scale bars are 1 mm. (c) Wild type; (d), 13U + e.v. and (e), 13U + *VTL8*, longitudinal sections of nodules stained with SYT013. Scale bars (c – e) are 200 μm.

In order to demonstrate which gene or genes caused the symbiotic phenotype of 13U, genetic complementation experiments were carried out. Transformation with constructs expressing *VTL4* and *VTL8* individually or together showed that expression of *VTL8* alone in the 13U mutant reverted the phenotypes to wild type. Specifically, nodules developed fully to nitrogen-fixing competency including the expression of leghaemoglobin and restoring the wild-type nodule zonation (Fig. 4b and e). Expression of *VTL4* alone in the 13U mutant did not have any effect, although co-expression with *VTL8* increased nodule fresh weight to above wild-type values (Fig. S2). Thus, the function of *VTL8* is essential for nodule maturation, whereas *VTL4, VTL5, VTL6* and *VTL7* only play a minor role if any.

### *MtVTL4* is required for early bacteroid development

The two *vtl4* mutant lines were indistinguishable from wild type with respect to shoot and root development (Fig. 5a). However, *vtl4* mutants had a relatively larger proportion of immature nodules compared to wild type (Fig. 5b). At 35 days post inoculation, wild-type plants had 20% immature nodules, compared to 34% and 57%, respectively, in *vtl4-1* and *vtl4-2* mutant plants. The number of immature nodules is more significant in the *vtl4-2* allele, in which the *Tnt1* insertion is closer to the start of the open reading frame. While no macroscopic differences were observed from the outside of the nodules (Fig. 5c), cross sections stained with SYTO13 showed that infected cells in zone II have a larger central vacuole and less elongated bacteroids (Fig. 5d). To quantify the difference, the diameter of the vacuoles of infected cells was measured in images of the differentiation zone of wild type and the *vtl4-1* mutant using ImageJ software. On average, the vacuoles in *vtl4-1* are 22.3 ± 5.7 μm (n = 284 cell images) and those in the R108 wild type 19.1 ± 4.0 µm (n = 88 cell images). This subtle cell morphological phenotype occurs in the zone where *VTL4* expression reaches its highest levels in nodules (Fig. 1b).

**Fig. 5.**
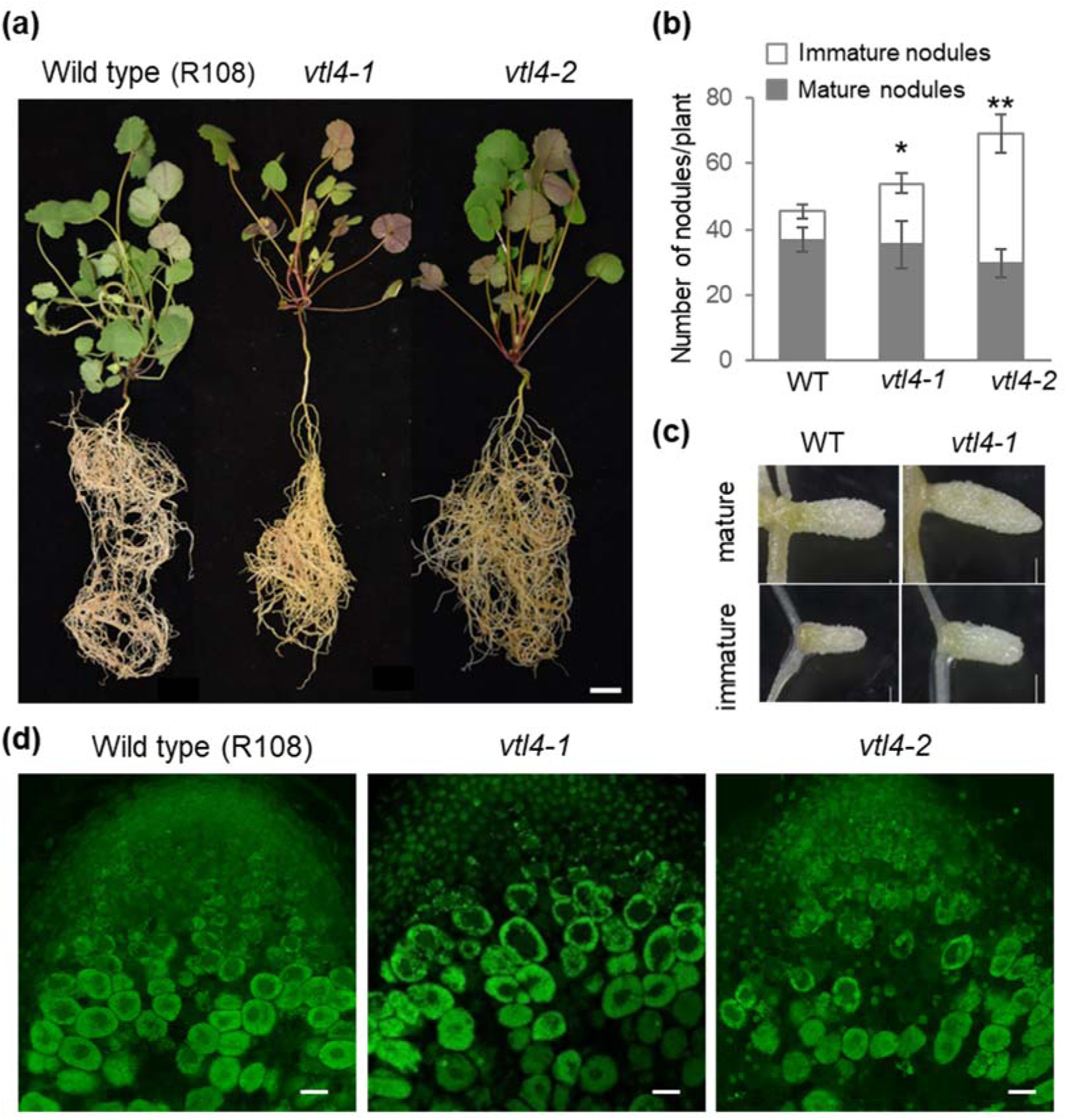
Disruption of *VTL4* delays infection. (a) *vtl4* insertion mutants and the parental wild type R108 inoculated with S. *meliloti* 1021, at 35 dpi. Scale bar is 1 cm. (b) The number of mature and immature nodules in *vtl4* mutants relative to wild type, at 35 dpi. *P<0.1, ** P<0.01 (n=6 plants), Student s t-test. (c) Representative images of mature and immature nodules in *vtl4*-1 and wild type. Scale bars are 1mm. (d) Longitudinal sections of nodules stained with SYT013, showing part of the infection zone (zone II). Scale bar is 100 μ m.

### Host plant VTLs are required for iron delivery to the bacteroids

It has previously been suggested that LjSEN1 and its homologs transport iron out of the infected plant cell into the peribacteroid space, based on sequence homology between VTLs and VIT proteins (Hakoyama *et al.*, 2011; Brear *et al.*, 2013; González-Guerrero *et al.*, 2016). In addition, Arabidopsis VTLs can complement the Δ*ccc1* yeast mutant defective in vacuolar iron transport, although growth complementation was only partial (Gollhofer *et al.*, 2014), and VIT proteins can transport manganese as well as iron (Lapinskas *et al.*, 1996).

To provide evidence for iron transport *in vivo*, we developed a transcriptional reporter in the bacteria using an iron-inducible promoter driving the *lux* operon. With the prerequisite that the bacterial promoter is active during nodule development, luminescence would be correlated with the iron status of the bacteria (deficient or sufficient). Most well-studied iron homeostasis genes, for example those for rhizobactin biosynthesis, are not expressed during nodule development and bacteroid differentiation (see Table S2 with data from Roux *et al.*, 2014). However, the expression of *mbfA* (membrane-bound ferritin A) is strongly induced in the proximal zone II and interzone, while its transcript levels are low in non-differentiated bacteria of Zone IId (Fig. 6a). The *mbfA* promoter is negatively regulated by the Iron-regulated repressor (Irr) which binds to the Iron Control Element (ICE) in the promoter (Rudolph *et al.*, 2006) (Fig. 6b). In addition to the wild-type promoter sequence of *mbfA*, we also made a reporter construct with a mutated ICE sequence, to prevent binding of Irr and achieve constitutive expression of *mbfA-lux*. Both reporter constructs were first tested in free-living *S. meliloti* 1021. Luminescence readings showed that the wild-type *mbfA* promoter was fully repressed under low iron but induced in the presence of iron (Fig. 6c). In contrast, the ICE mutant promoter construct (*PmbfA*^ICE^:*lux*) was de-repressed in the absence of iron, although some Fe-dependent regulation was still observed.

**Fig. 6.**
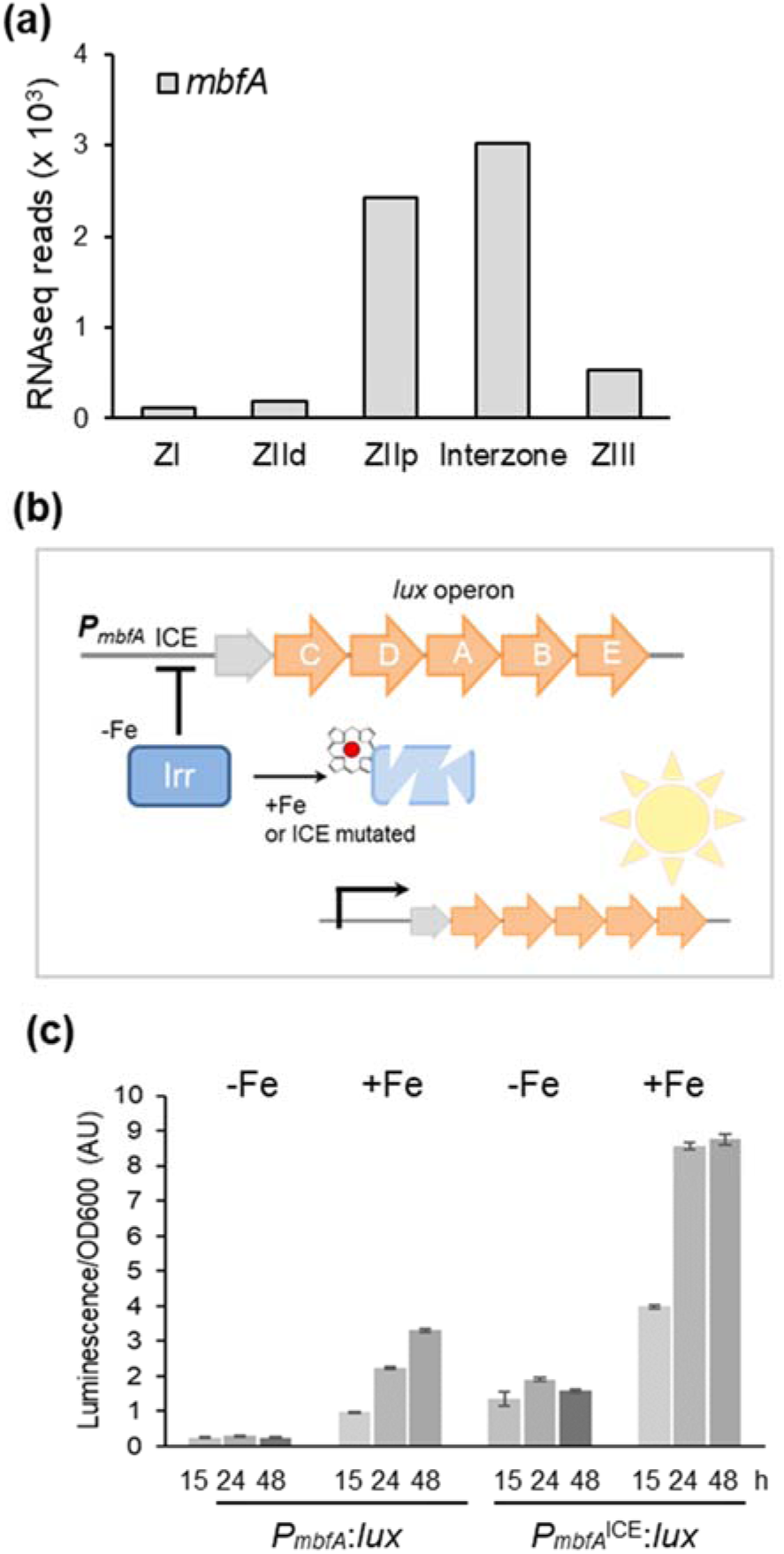
Development of an iron reporter for symbiotic S. *meliloti.* (a) Expression levels of S. *meliloti mbfA* during nodule development. Data were obtained from RNAseq data published by Roux et al. (2014). (b) Diagram of the bacterial P*_mbfA_*:*lux* reporter construct, including the Iron-regulated repressor Irr and the Iron Control Element (ICE). (c) Expression of P*_mbfA_*:*lux*, measured as luminescence, in minimal medium without iron (-Fe) or supplemented with 80 pM iron sulphate (+Fe). Values are the average luminescence readings corrected for OD_600_ of 3 replicate cultures. Error bars represent ± SD.

Next, wild-type and mutant *M. truncatula* seedlings were inoculated with bacteria expressing the *mbfA*-*lux* reporter, and luminescence was analysed at 21 dpi using a NightOwl imaging system. *S. meliloti* 1021 was used to inoculate all plant genotypes. While less effective in nodulating *M. truncatula* Jemalong, the parent line of the 13U mutant, the number of nodules were sufficient for quantitative analysis. Luminescence was only observed in nodules, where bacteria are concentrated and provided with iron (Fig. 7a). Quantification of the signal showed approximately 50% luminescence in nodules of *vtl4* mutants, and only 10% in the 13U mutant, compared to their respective wild types (Fig. 7b). Using the ICE mutant reporter, the luminescence signal was similar in the *vtl4* mutants and parental line. In the 13U line, luminescence was slightly increased compared to the control line (Fig. 7c). Thus, the number of bacteria in Zone II and the Interzone in wild-type and mutant nodules are comparable, regardless of their difference in nodule development. Taken together, the lower expression of the bacterial iron reporter in the *M. truncatula vtl4* and 13U mutants indicates that the nodule-specific VTL proteins are required for iron delivery to the bacteroids.

**Fig. 7.**
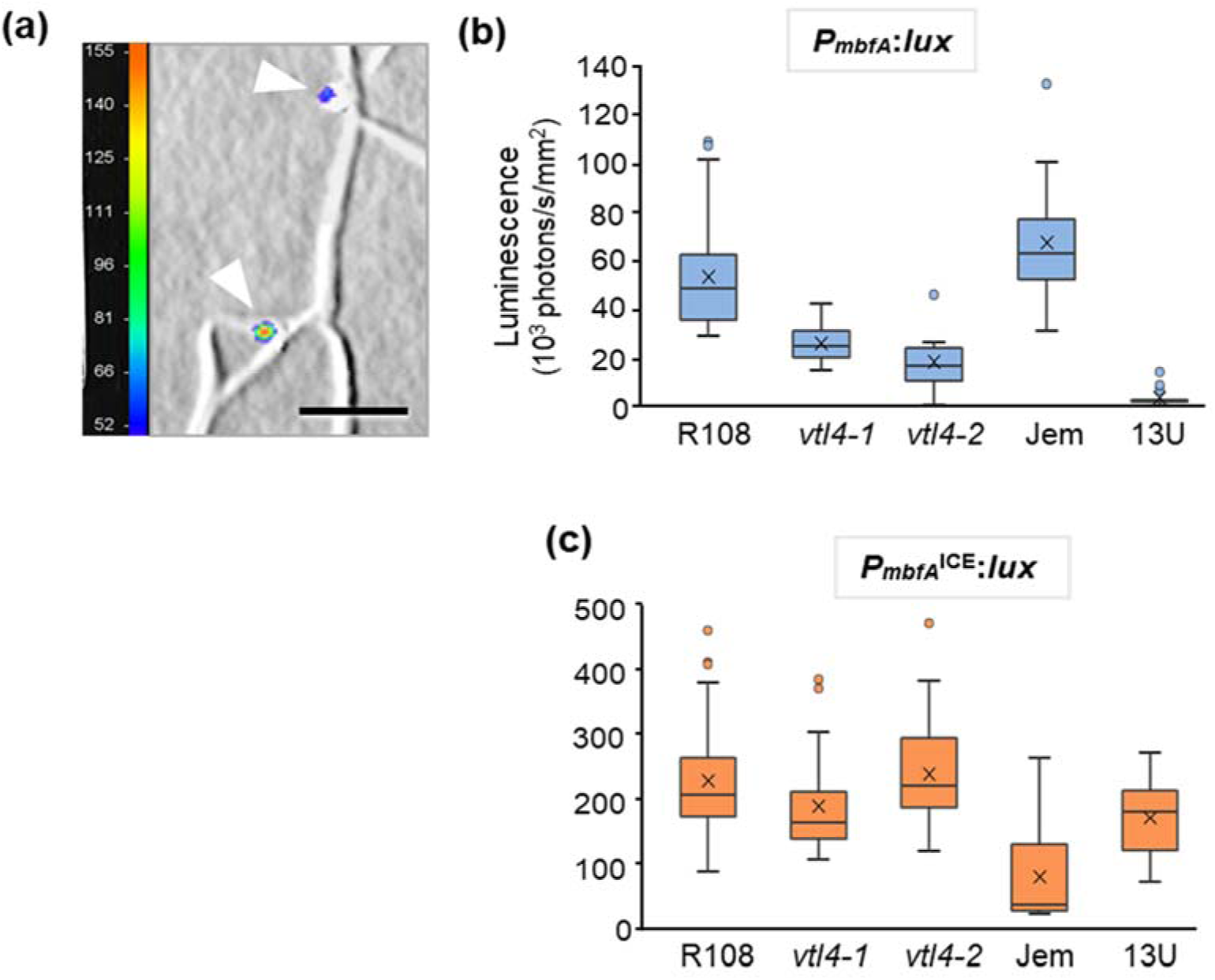
Symbiotic bacteria in nodules lacking VTL4 and VTL8 are iron deficient. (a) Detail of an *M. truncatula* R108 root inoculated with S. *meliloti* 1021 expressing P*_mbfA_*:*lux* at 21 dpi. Luminescence is represented by a colour scale in photons per second, superimposed on a grayscale image. The white arrow heads point to nodules. Scale bar is 5 mm. (b, c) Expression of the bacterial Fe reporter P*_mbfA_*:lux (b) and the deregulated ICE mutant (c) in root nodules at 21 dpi, measured as luminescence using a NightOwl camera. *M truncatula* R108 was used as wild type for the *vtl4* mutants, and *M. truncatula* Jemalong (Jem) as wild type for the 13U mutant. The data are presented in a box plot, with the box marking the upper and lower quartiles, the middle line represents the median and the symbol x indicates the mean. The whiskers show the spread of the data within the 1 5-fold interquartile range (n=9 for *Vtl4-2*, and n≥13 for other lines).

### VTL4 and VTL8 mediate iron transport into yeast vacuoles

To confirm that VTL4 and VTL8 are able to transport iron, the genes were expressed in yeast for functional complementation assays. The *VTL* genes were cloned into the pYES2 plasmid under the control of a galactose-inducible promoter. The plasmids were transformed into a Δ*ccc1* yeast mutant which lacks the vacuolar iron transporter Ccc1 (Li *et al.*, 2001). Its inability to store iron in the vacuole leads to severely impaired growth in the presence of excess iron in the medium (Fig. 8). Growth can be restored by expressing another vacuolar iron transporter such as Arabidopsis *VIT1* (Kim *et al.*, 2006), used here as a positive control alongside the wild-type strain (Fig. 8). *VTL4* as well as *VTL8* were able to rescue growth of the Δ*ccc1* strain on medium with 5 mM Fe sulphate. Functional complementation was only seen using Δ*ccc1* in the DY150 (W303 derivative) genetic background, but not in the BY4741 strain. This may be due to differences in salt tolerance between the two strains causing an indirect effect on Fe homeostasis (Petrezxelyova *et al.*, 2010).

**Fig. 8.**
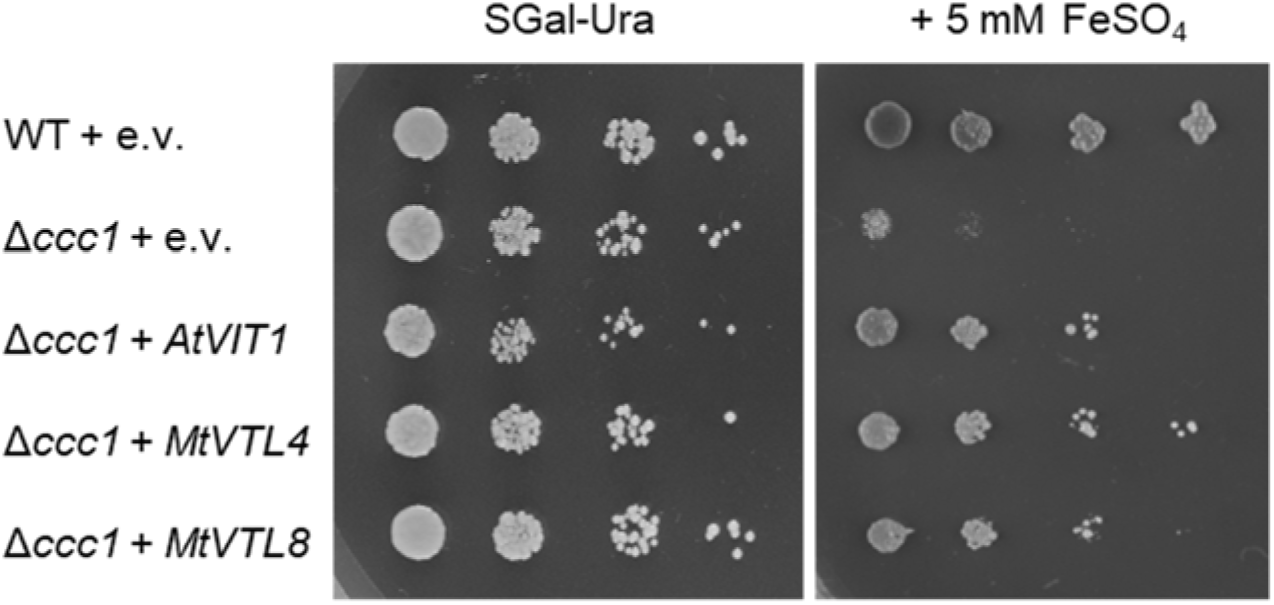
Expression of *VTL4* or *VTL8* restores vacuolar iron transport in Δ*ccc1* yeast. Yeast transformed with pYES2 empty vector (e.v.) or with the indicated gene were grown on minimal synthetic medium with galactose (SGal) lacking uracil (-Ura) in a 5-fold serial dilution. Addition of 5 mM iron sulfate to the same medium selects for strains able to transport iron across the vacuolar membrane.

## DISCUSSION

Because of their high demand for metals, nodules provide an interesting system to study the function of transporters and other metal homeostasis genes in a highly compartmentalized plant cell. After iron is imported across the plasma membrane into the infected cell, it is distributed between the mitochondria, plastids containing the iron storage protein ferritin, and the differentiating bacteroids. Because of toxicity of ‘free’ iron (not bound to chelators or proteins), the timing of distribution and thus gene expression is crucial. In *M. truncatula*, 2 of its 8 *VTL* genes appear to be involved in iron delivery to the bacteroids, but at different times of the infection process. This may reflect the findings of older studies in peanut and lupin, that iron is required for both nodule initiation and further development (O’Hara *et al.*, 1988; Tang *et al.*, 1990). VTL4 is providing iron to the bacteria, before any differentiation, when they are dividing in the infection thread. The minor effects of knocking out *VTL4* on nodule and bacteroid development suggest that VTL4 is not the only source of iron. At this stage, the bacteria may still have some iron stored from before infection, or simply need little iron. In contrast, VTL8 is critical for full differentiation of the bacteroids into nitrogen-fixing symbionts. In line with this, *VTL8* expression peaks just before the induction of the bacterial *nif* genes for nitrogenase when large amounts of iron need to reach the bacteroids for incorporation in the iron-sulphur cofactors of the abundant nitrogenase protein.

Phylogenetic analysis shows that *VTL8, LjSEN1* and *GmNOD21* form a separate clade of *VTL* genes. It is therefore likely that the gene was recruited specifically for symbiosis in the ancestral legume species. Why was a *VIT-like* gene, but not *VIT*, co-opted for this function? VIT is a Fe^2+^/H^+^ antiporter, but the iron species transported by VTLs is not known. Because the iron-binding loop is missing in VTLs, it is possible VTLs transport both Fe^2+^ and Fe^3+^, or Fe^3+^ exclusively. Interestingly, VTLs are highly homologous to the transmembrane domain of MbfA, which also lacks the Fe-binding loop but has an additional N-terminal ferritin-like domain (Bhubhanil *et al.*, 2014; Sankari & O’Brian, 2014). It has been suggested that this domain oxidizes Fe^2+^ to Fe^3+^, which is then transported by the membrane domain. The ferritin-like domain may also help drive transport in one direction, namely to the outside of the cell. It would be interesting to see if plant ferritin, which accumulates transiently in Zone II, is able to associate with VTL8 to deliver iron to the peribacteroid space. Normally the 24-mer iron-storage protein ferritin is localized on the periphery of plastids in plants (Roschzttardtz *et al.*, 2013; Moore *et al.*, 2018). So, either plastids interact with the symbiosome membrane, or a cytosolic form of ferritin exists in infected cells.

It is still not clear how iron exported from the plant cell into the peribacteroid space is taken up by the bacteroids. The peribacteroid space undergoes acidification towards the onset of nitrogen fixation as shown by fluorescent probes (Pierre *et al.*, 2013). The pH is a critical factor for the solubility of iron, with a low pH enhancing solubility. Most bacteria species have separate transport systems for Fe^2+^ and Fe^3+^-chelate complexes. Iron uptake genes that are active in free-living rhizobium (Johnston *et al.*, 2001) are generally not induced in the symbiont stage (Table S2, Roux *et al*., 2014, and reviewed in (Abreu *et al.*, 2019). The Fe^2+^ transporter FeoAB plays a role in iron transport in *Bradyrhizobium* and *feoA* or *feoB* deletion strains produced ineffective nodules on soybean (Sankari & O’Brian, 2016). However, there are no *feoAB* genes in the *S. meliloti* genome. Alternatively, iron may be taken up as Fe^3+^-citrate of Fe^3+^-malate via tricarboxylic acid transporters, which are highly induced. The Fe-responsive *lux* reporter developed for this study should help to fully characterize the bacterial iron transporters that are active in the nodules. It is important to note that the strongly induced *MbfA* expression indicates bacteroids are saturated with iron, and need to export it back to the peribacteroid space. Thus, low-affinity uptake systems may be sufficient for the bacteroids’ Fe requirements.

In summary, this study provides confirmation that VTL8 / SEN1 is indeed the main iron transporter across the plant symbiosome membrane, but it also opens up new questions regarding the iron homeostasis in nodules. In addition to identifying the Fe-species that is exported, keys questions are how Fe is delivered to VTL8 / SEN1 and how Fe is partitioned between haem biosynthesis for leghaemoglobin and export to the bacteroids.

## MATERIALS AND METHODS

### Plant growth and bacterial strains

*Medicago truncatula* Jemalong and *M. truncatula* subsp. *tricycla* (R108) genotypes were used as wild-type controls for the 13U (Domonkos *et al.*, 2013) and *vtl4* mutant lines, respectively. The *Tnt1*-insertion mutants *vtl4-1* (NF17463) and *vtl4-2* (NF21016) were purchased from the Samuel Roberts Noble Foundation (Ardmore, Oklahoma). *M. truncatula* seeds were scarified with sandpaper before being surface sterilised with 10% (w/v) sodium hypochlorite for 4 min. Seeds were then washed 5 times with distilled sterile water and imbibed in the dark at room temperature for 4 h. Seeds were plated on sterile water agar, and stratified by placing the plates upside down at 4 °C for 3-7 days. The plates were then moved to 20 °C for 16 h to allow the seeds to germinate. Seedlings were transferred either to sterile medium or planted out on a 50:50 mixture of Terragreen and sand for inoculation with rhizobium. Plants were grown in long day conditions (16 h light, 8 h dark) at 22 °C, light intensity 180-200 μmol photons m^−2^ s^−1^.

Two different bacterial strains were used, as indicated in the figure legends. *Sinorhizobium* (*Ensifer*) *medicae* WSM419 (pXLGD4 *PhemA*:*lacZ, tetR*) was used for nodulating *M. truncatula* Jemalong and the 13U mutant; *Sinorhizobium meliloti* 1021 (*PnifA:lacZ, tetR*) was used for *M. truncatula* subsp. *tricycla* (R108) and the *vtl4* mutants. *S. meliloti* 1021 can also establish nitrogen-fixing symbiotic interaction with Jemalong albeit less effectively, and was used in the *lux* reporter assays for all plant lines. Bacteria were grown in TY medium (1% (w/v) tryptone, 0.3% (w/v) yeast extract, 6 mM CaCl2), either in liquid or solid medium with 1% (w/v) agar.

### Identification of the 13U mutation and genotyping of *vtl4* lines

A combined approach of genetic mapping and microarray-based Affymetrix GeneChip hybridization was applied to identify the deletion in the 13U mutant line which was isolated from a collection of fast neutron radiation mutants. Initial analysis had placed 13U in linkage group 4 of *M. truncatula* (Domonkos *et al.*, 2013). The 13U mutant was crossed with the A20 genotype and more than 250 F2 individuals were scored for symbiotic phenotypes in relation to the genetic markers amplified with the primer pairs listed in Table S1. To identify deletions in the genome of 13U, genomic DNA from the 13U mutant was labeled and hybridized to an Affymetrix GeneChip, as described previously (Murray *et al.*, 2011). To determine the boundaries of the deletion on chromosome 4, the region was scanned by PCR amplification using primers designed for the predicted genes in the region (Table S1). For genotyping *vtl4* mutants, genomic DNA was used as template for PCR reactions with one primer set designed to amplify the wild-type *VTL4* gene and a second primer set with one primer in the *VTL4* coding sequence and the other in the *Tnt1* sequence to detect the insertion.

### Gene expression analysis

To assess *VTL4* and *VTL8* transcript levels, total RNA was extracted from wild-type and mutant nodules using the RNeasy Mini kit (QIAGEN, Germany) and treated with DNase (Turbo DNase kit, Agilent). cDNA was produced using Thermo SuperScript II Reverse Transcriptase and an anchored oligo-dT primer, and used as template for either standard RT-PCR with products separated by agarose gel electrophoresis, or for quantitative RT-qPCR. RT-qPCR reactions were made using SensiFAST master-mix (Bioline), each with 20 ng of cDNA. Reactions were measured in a Bio-Rad CFX-96 real-time PCR system and cycled as per the Bioline protocol. The expression values were normalized to that of the *UPL7* gene (*UBIQUITIN PROTEIN LIGASE 7, Medtr7g103210.1*)

### *Agrobacterium rhizogenes*-mediated complementation of *M. truncatula* mutants

Fragments containing the genes *VTL4* and *VTL8* including the 1.6 and 1.5 kb native promoter sequences and 1.2 and 0.9 kb 3’untranslated regions, respectively, were amplified with the oligonucleotides listed in Table S1 and using BAC (Bacterial Artifical Chromosome) clone mth2-28d20 as a template. The destination vector pKGW-R (gateway.psb.ugent.be/vector/show/pKGW_RedRoot/) was linearized by digestion with restriction enzymes *Aat*II and *Spe*I. The PCR-generated gene fragments and the linearized vector were fused using the In-Fusion HD Cloning Kit (Thermo Fisher Scientific) according to the manufacturer’s protocol. Constructs were introduced into ARqua1 strain of *A. rhizogenes* and used for hairy root transformation as described in the *Medicago truncatula* Handbook (www.noble.org/MedicagoHandbook).

### Fluorescence microscopic analysis of nodules

Transformed roots were identified by fluorescence of the DsRed marker protein. To analyze nodule occupancy by rhizobia, nodule sections were stained with the DNA-binding fluorescent dye SYTO13 (Invitrogen, Eugene, Oregon). The preparation of nodule sections was carried out as described earlier (Domonkos *et al.*, 2013). Sections were stained in 1×PBS (pH 7.4) containing 5 μM SYTO13 for 20 min and rinsed with 1×PBS before analysis. Images were acquired with a Leica TCS SP8 confocal laser scanning microscope with the following configuration: objective lens: HCX PL FLUOTAR 10x/0.30 (dry, NA:0.3), PL FLUOTAR 40x/1.00 OIL and HC PL APO CS2 63x/1.40 OIL, scanning mode: sequential unidirectional; excitation: 488 nm; main dichroic beamsplitter: DM488/552; detection range for the SYTO13 channel was 507-564 nm.

### Protein localization

The *VTL4* promoter (2566 nt), *VTL8* promoter (2096 nt) and coding sequences were domesticated for Golden Gate assembly. A glycine-rich linker sequence (Table S1) was added to the C-terminus followed by the coding sequence of mCherry. As a marker for the plasma membrane, the *A. thaliana PIP2A* (*AT3G53420*) sequence was fused to eGFP and placed behind the promoter of *L. japonicus UBIQUITIN* (GenBank DQ249171.1) The sequences were assembled into backbone vector pL1-R1 and transformed into *Agrobacterium rhizogenes* strain ARqua1. *M. truncatula* seeds were sterilized and germinated on water-agar containing 3 mg/ml Nystatin. Approximately a quarter of the root of germinated seedlings was removed before dipping in a suspension of *A. rhizogenes* bearing the plasmid of interest. Seedlings were then plated out on modified Fahraeus medium (0.7 mM KPO_4_, 0.8 mM Na_2_PO_4_, 0.50 mM MgSO_4_, 0.7 mM CaCl_2_, 20 µM Fe-citrate, pH 6.0, and micronutrients (5 µM H_3_BO_3_, 9 µM MnSO_4_, 0.8 µM ZnSO_4_, 0.3 µM CuSO_4_ and 0.5 µM H_2_MoO_4_) with Nystatin. Plants were grown at a 45° angle in the dark for a week before transferring to fresh plates lined with filter paper in the light. After 3 weeks plants were transferred to fresh modified Fahraeus plates containing Nystatin and 100 nM aminoethoxyvinylglycine (AVG) and inoculated with 0.5 mL *S. meliloti* 1021 at an OD600 of 0.03 and grown for 21 days. Nodules were harvested, hand-bisected and mounted on microscope slides on double-sided tape, with water as the mounting medium and hi-vac silicone grease as a spacer. Fluorescence was imaged using a Leica TCS SP5 confocal microscope. For eGFP detection the excitation wavelength was 488 nm, emission was measured at 493 – 550 nm with a Hybrid (HyD) detector; for mCherry detection the excitation wavelength was 561 nm, emission was measured at 575 – 650 nm; and for DAPI (DAPI, or 4′,6-diamidino-2-phenylindole) the excitation wavelength was 405 nm, emission was measured at 435 – 477 nm.

### *PmbfAp:lux* reporter and expression analysis in *S. meliloti* 1021

Nucleotides −487 to +104, containing the promoter sequence and part of the coding sequence of the *mbfA* gene (SMc00359) were PCR-amplified from *S. meliloti* 1021 (see Table S1 for primer sequences) and cloned using the *Bam*HI and *Kpn*I restriction sites into pIJ11268 upstream of the *lux* operon (Frederix *et al.*, 2014). A de-repressed version of the reporter was made by mutating the Iron Control Element (ICE) from TTCTAA to AGCTTC (−19 to −14) by site-directed mutagenesis. The plasmids were transferred from *Escherichia coli* to *S. meliloti* by conjugation using a helper strain carrying plasmid pRK2013. For luminescence assays, overnight bacterial cultures were diluted in UMS medium (Wheatley *et al.*, 2017) containing 20 mM 3-morpholinopropane-1-sulfonic acid-KOH pH 7, 10 mM glucose, 10 mM NH_4_Cl, 8.5 mM NaCl, 2 mM MgSO_4_, 0.5 mM K_2_HPO_4_, 0.51 mM CaCl_2_, 1x Trace Elements (1 µM Na_2_-EDTA, 0.6 µM ZnSO_4_, 0.8 µM Na_2_MoO_4_, 4 µM H_3_BO_3_, 0.9 µM MnSO_4_, 0.08 µM CuSO_4_, 4 µM CoCl_2_, 3 µM thiamine, 4.2 µM D-pantothenic acid, 0.4 µM biotin, without or with 80 µM FeSO_4_ as indicated. Luminescence and OD600 of triplicate cultures for each strain and condition were measured in a CLARIOstar microplate reader (BMG LABTECH).

### Luminescence imaging and quantification

Plants were inoculated with *S. meliloti* 1021 carrying the *PmbfA:lux* reporter plasmid or the mutant form *PmbfA*^ICE^:*lux* at 7 days postgermination, and grown for a further 21 days. Plants were dug up, roots rinsed and blotted dry and imaged using the NightOWL II LB 983 in-vivo imaging system with IndiGO software (Berthold Technologies, Bad Wildbad, Germany). Luminescent nodules were identified using the automated peak picking tool and the luminescence in photons per mm^2^ calculated from the output. Five plants per line were analysed with each inoculum.

### Yeast complementation

The yeast strain DY150, which is derived from W303, was used as wild type. The Δ*ccc1* strain in this background carries a genomic deletion of *CCC1*, initially identified as *Cross-Complements Ca*^*2+*^*1*, but later shown to mediate vacuolar iron transport (Li *et al.*, 2001). Plant genes were cloned into shuttle vector pYES2 under the control of the *GAL1* promoter for galactose-inducible expression. The coding sequence of Arabidopsis *VIT1* (*AT2G01770*) was used as a positive control for functional complementation. Yeast were transformed using the lithium-acetate method and positive transformants were selected on synthetic dropout medium lacking uracil (DSCK102, Formedium, Hunstanton UK) with glucose as carbon source (SD). Overnight cultures of selected colonies were grown in SD-Ura, then spotted onto 2% (w/v) agar plates of SGal-Ura with or without 5 mM FeSO_4_. Plates were photographed after 4 days (control) or 5 days (with iron).

## Acknowledgements

We thank Jeremy Murray, Andy Breakspear and Giles Oldroyd for their generous advice to J.H.W.’s PhD research and for providing gene sequences for Golden Gate cloning. We also like to thank Jeremy Murray for help with identification of the deleted genome region in the 13U line; Allan Downie (John Innes Centre) for the *lux* plasmid; Penelope Smith (La Trobe University) for sending *AtVIT1* and *MtVTL4* in pYES2; Thomas Buckhout (Humboldt University Berlin) for the Δ*ccc1* and wild-type yeast strains; Eva Wegel (John Innes Centre) and Zoltán Tóth (Agricultural Biotechnology Institute) for confocal microscopy; Hannah Justice and Heather Bland for phenotype analysis. We thank Kristina Miró (ABC, Gödöllő) for skillful technical assistance. This study was supported by Biotechnology and Biological Sciences Research Council (BBSRC) Institute Strategic Grants BB/J004553/1 and BB/P012574/1 (R.G.T., M.F. and J.B.), the Gatsby Charitable Foundation (J.H.W.), and the Hungarian Scientific Research Fund grants NKFIH OTKA 67576 and 119652 (P.K.).

## Author Contributions

J.H.W., R.T.G., G.K.K., A.D., B.H., E.M.B. and M.F. performed the research and helped with designing experiments and data analysis; P.K. and J.B. analysed and interpreted the data and wrote the manuscript.

