## Supplementary material for "The *Medicago truncatula* Vacuolar Iron Transporter-Like proteins VTL4 and VTL8 deliver iron to endosymbiotic bacteria at different stages of the infection process": Suppl Figs 1 and 2

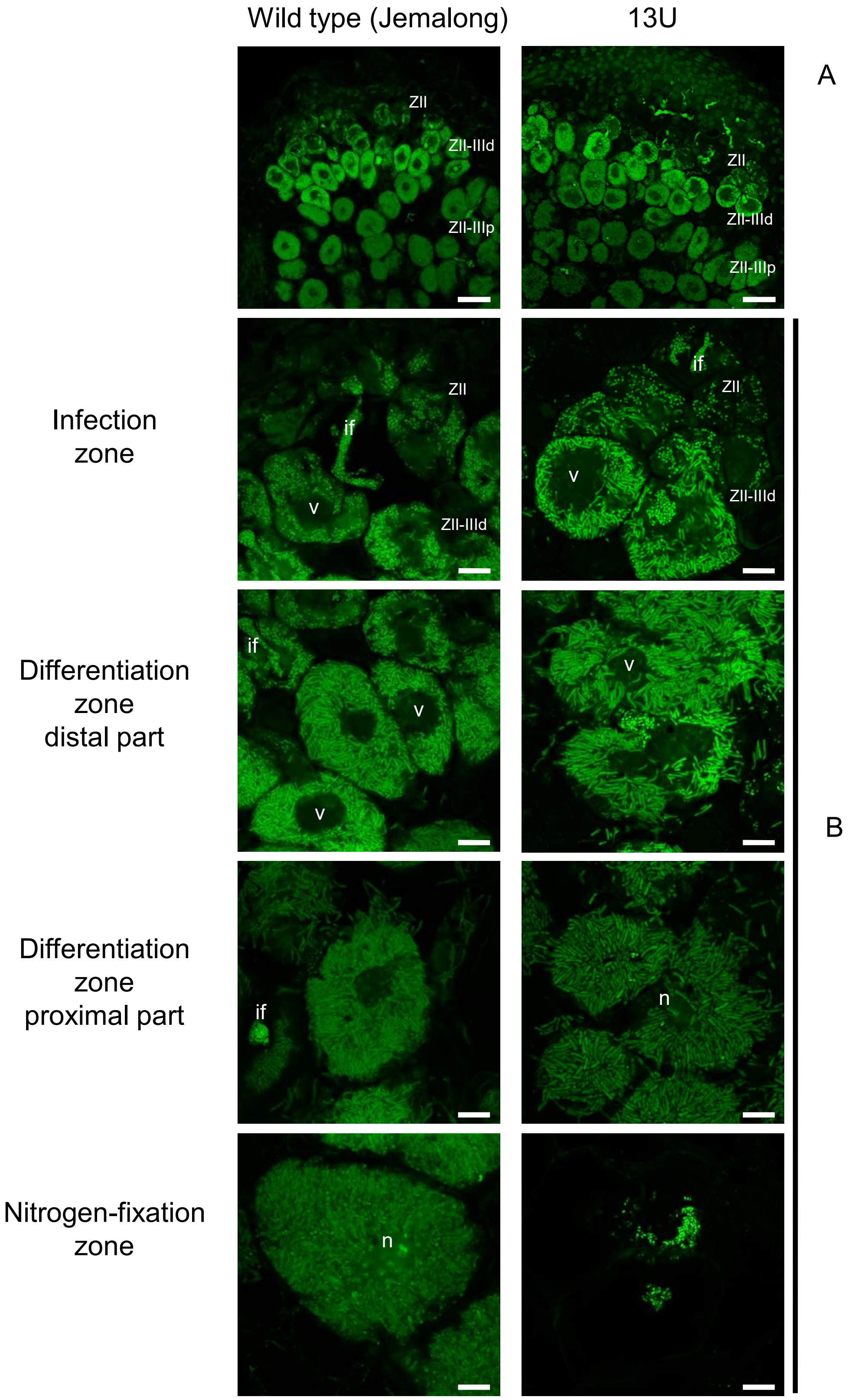


**Supplemental Figure S1**. Cytology of infected cells in the *M. truncatula* 13U mutant. Confocal images of SYTO13-stained wild-type (Jemalong) and 13U nodule sections three weeks post inoculation with *S. medicae* WSM419. 13U nodules show disorganized dispositon of elongated bacteroids in the differentiation zone and have a nitrogen-fixation zone devoid of symbiotic cells. Scale bars (A) 50 μm, (B) 10 μm. ZII: infection zone, ZII-IIId: differentiation zone distal part, ZII-IIIp: differentiation zone proximal part. if: infection thread, v: vacuole, n: nucleus.

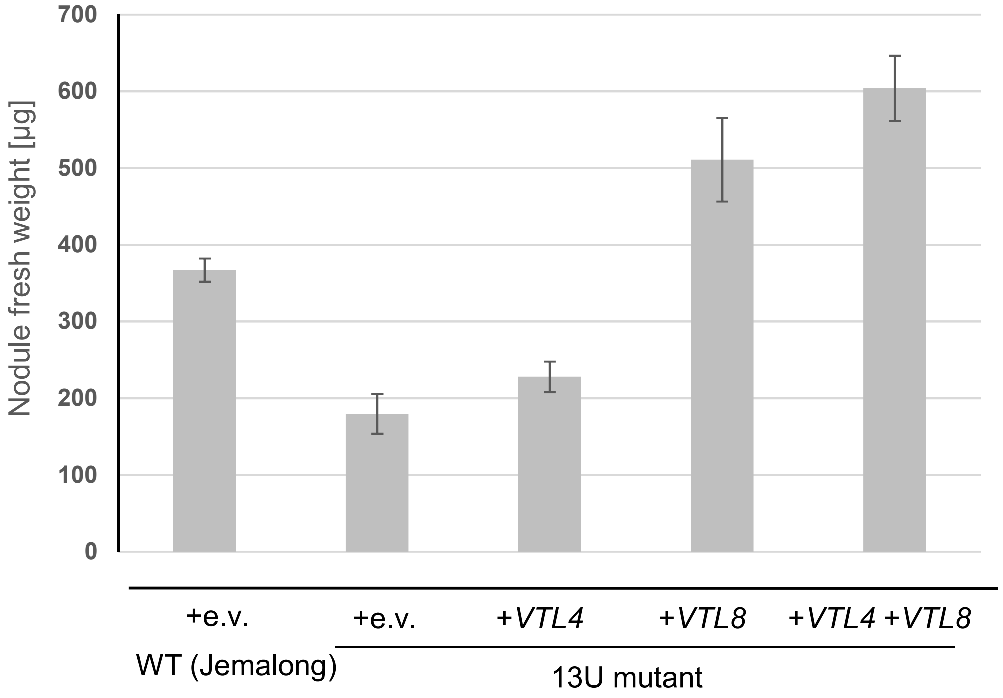


**Supplemental Fig. S2**. Nodule fresh weight of complemented 13U lines. For each root transformation, nodules were pooled to determine the total fresh weight, divided by the number of nodules. The values are the mean of 3 independent transformations ± SE.
