## Supplementary material for "The *Medicago truncatula* Vacuolar Iron Transporter-Like proteins VTL4 and VTL8 deliver iron to endosymbiotic bacteria at different stages of the infection process": Suppl Tables 1 and 2

**Supplemental Table S1. Primers and other oligonucleotides**

| **Oligonucleotide** | **Purpose** | **Sequence (5’-3’)** |
| --- | --- | --- |
|  | Genotyping of Tnt lines | TCCTTGTTGGATTGGTAGCC  CAGTGAACGAGCAGAACCTGTG |
| VTL4F (JW65)  VTL4R (JW83) | Genotyping of *vtl4* mutants | CAACCTCCAACCACCACTCT  TGTACTTCATGGCTTCTCTTGG |
| JW3  JW4 | RT-PCR of *VTL4* | AGCTGCTTTGTTAGGAGCCA  ATGTCTTCATGCACAGCTCCA |
| VTL4RTqF  VTL4RTqR | RT-qPCR of *VTL4* | ACCACCAGAAATGAGAAAGAGC  TTGAAGTTGGCCACCTCTGA |
| SEN1qRTF  SEN1qRTR | RT-PCR and qPCR of *VTL8* | GCCATAGCCATCCATCCACC  GGCTTTGGTTGTGTTTGGAGG |
| UPL7qRTF  UPL7qRTR | RT-qPCR reference gene | CCAGTTGTTCTCGTGGTCCATT  CCTCCAATTGTCGCCCAAA |
| FL3 | Linker peptide  LGGGGSGGGGSGGGGSAAA | CTTGGTGGAGGTGGATCTGGTGGAGGTGG  ATCAGGTGGAGGTGGATCTGCTGCAGCT |
|  | Cloning of *MtVTL4* into pYES2  (Two-step PCR for Gateway) | AAAAAGCAGGCTCCATGGCTTCTCTTGGTAACCATAAT  AGAAAGCTGGGTATCAAAGTGAACTATAGCCAACGAA |
| M39  M40 | Cloning of *MtVTL8* into pYES2  (One-step PCR for Gateway) | ggggacaagtttgtacaaaaaagcaggctcgATGGCCGTTGGTACAATATG  ggggaccactttgtacaagaaagctgggtaTCAAATTTCCAAATCCAAACC |
|  | Cloning of *AtVIT1* into pYES2  (One-step PCR for Gateway) | ggggacaagtttgtacaaaaaagcaggcttaATGTCGTCGGAGGAAGATAAGA  ggggaccactttgtacaagaaagctgggtaCTAATGTTGCACAACTTTAGCC |
| mbfANEWF  mbfANEWR | Cloning of *S. meliloti* *mbfA* promoter | ACTGGGTACCGACAAGGTGGTGGCGATGTA  ACTGGGATCCAATGCGTCCATCGTCCTCTTC |
| ICESDM2F  ICESDM2R | Mutagenesis of *mbfA* ICE box | CATCACGAAGCCTTCTGAAGCTTAATTCTAATTAGGCGCCGATTCATGACGCG  CGCGTCATGAATCGGCGCCTAATTAGAATTAAGCTTCAGAAGGCTTCGTGATG |
| KGWSpeI_325pr_F  KGWAatII_325UTR_R | Cloning of *VTL4* into vector pKGW-R | aggcggccgcactagTCGTGACGGAAAACATGTAGGG  gcaccaaccacaacgacgtTGAGCTGAATTTTTGAGCTGACTATAC |
| KGWSpeI_335pr_F KGWAatII_335UTR_R | Cloning of *VTL8* into vector pKGW-R | aggcggccgcactagCCAAACGACATGGCATATAGC  gcaccaaccacaacgacgtCATATAAACCTCCCGAAACCCTC |
| 325prom_F  335_R&325prom_hom | Cloning of *VTL8 + VTL4* into vector pKGW-R | TCGTGACGGAAAACATGTAGGG  TGTTTTCCGTCACGACATATAAACCTCCCGAAACCCTC |
| h2_16e5_F  h2_16e5_R  h2_17n4b_F  h2_17n4b_R  h2_68o3b_F  h2_68o3b_R  mth2-49e15_F  mth2-49e15_R mth2_72d18e_F  mth2_72d18e_R | Fine mapping of 13U | CTGCAATTAAACAAAATGGTTAGAGT  CTACCAAACAAAAGCCTAGACTTCTT  CAACGCCGGGTAAATACACT  TTGTTTTTGTGTTTCGCACA  GGTTCAACAATCAAATGAGCTG  GTGAGCAGAAGCATACACGC  AACCCCAGACACCCCACT  GCACGCATGCACACTCAT  TGACCAATGAAGCGCAATAC  TATTAAAACATCCCGACCGC |
| BG646574_F2  BG646574_R2  SYN24_F  SYN24_R  TC173960_F1  TC173960_R1b  TC178276_F1  TC178276_R1  TC182270_F1  TC182270_R1 | Delimit the 13U deletion | GAGTCAAACGAAGTAAAGGTT  ACTCAAAACTGGAACTCTGAC  TCAACTCAACAATGCTTACTATGGC  AAAGAGGAAACCAATGATTGCTACT  TCACCATTGCCCCACTGATGCTA  ACAACCTTATTCCAGGAGTTACC  GGCAAAGGGCTCAATGGCTTCGC  GCACCAACACCTCCAAACACAAC  TTCTTTGCCTTTTTATTTTTCGTTC  TGGTTCAGGTCCTTCATTAGTCTTG |
| TC192817_F1  TC192817_R1  CA919468_F1  CA919468_R1  TC191603_F1  TC191603_R1 | Detecting borders of the 13U deletion | CAACAAACCAAAAGACCAGCATAGG  AGGGATGGTGGTAGTCGTGGAGATT  AATGAATGACCCTATCTCACGGA  ATTTCTACATCTGCACTTGGCTT  TCAGGCTTCTCACTATGGCTTATT  CACTGACCCAGAGACCACAATACC |

**Supplemental Table S2. Expression of *S. meliloti* genes involved in Fe homeostasis during nodule development**

**Data from Roux *et al*., (2014)** An integrated analysis of plant and bacterial gene expression in symbiotic root nodules using laser-capture microdissection coupled to RNA sequencing. Plant J 77: 817–837.

|  | | Total reads | Percentage (%) expression | | | | |
| --- | --- | --- | --- | --- | --- | --- | --- |
| ***M. truncatula*** | |  | F1 (zone I) | FIId | FIIp | IZ | ZIII |
| *VTL4* | Medtr4g094325 | 40662.4 | 3.9 | 44.1 | 8.6 | 16.7 | 26.8 |
| *VTL8* | Medtr4g094335 | 150938.1 | 0.1 | 0.2 | 7.3 | 68.6 | 23.8 |
| ***S. meliloti*** | |  |  |  |  |  |  |
| *bfr* (putative bacterioferritin) | SMc03786 | 6959.4 | 21.6 | 24.6 | 22.7 | 26.2 | 4.9 |
| *fecI* (Fe citrate transport, regulation) | SMc04203 | 188.5 | 13.8 | 16.5 | 32.2 | 21.2 | 16.3 |
| *fecR* | SMc04204 | 643.0 | 2.0 | 11.4 | 40.6 | 24.3 | 21.7 |
| *hmuV* (hemin uptake) | SMc01510 | 556.0 | 20.5 | 30.5 | 25.3 | 11.0 | 12.9 |
| *hmuU* | SMc01511 | 1240.2 | 24.3 | 26.1 | 27.8 | 11.9 | 9.9 |
| *hmuT* | SMc01512 | 1655.5 | 25.4 | 22.1 | 33.5 | 14.2 | 4.9 |
| *hmuS* | SMc01513 | 3746.6 | 32.6 | 18.4 | 29.9 | 15.3 | 3.8 |
| *mbfA* (membrane-bound ferritin) | SMc00359 | 6289.4 | 1.9 | 3.0 | 38.7 | 48.0 | 8.4 |
| *rhbA* (rhizobactin biosynthesis) | SMa2400 | 391.5 | 70.0 | 4.6 | 5.3 | 5.6 | 14.5 |
| *rhbB* | SMa2402 | 391.5 | 70.0 | 4.6 | 5.3 | 5.6 | 14.5 |
| *rhbC* | SMa2404 | 961.7 | 32.3 | 17.5 | 25.2 | 15.1 | 10.0 |
| *rhbD* | SMa2406 | 515.2 | 49.8 | 11.6 | 8.6 | 12.2 | 17.8 |
| *rhbE* | SMa2408 | 131.6 | 66.0 | 7.2 | 8.0 | 7.9 | 10.8 |
| *rhbF* | SMa2410 | 955.8 | 26.4 | 14.7 | 14.9 | 8.9 | 35.1 |
| *rhtA (rhizobactin transporter)* | SMa2414 | 3406.1 | 65.0 | 10.6 | 4.9 | 6.2 | 13.3 |
| *Putative Fe/heme transporter* | SMc04205 | 461.9 | 11.1 | 8.6 | 16.0 | 23.4 | 41.0 |
